# An optimized RNA polymerase II minigenome system for Nipah virus

**DOI:** 10.64898/2026.06.12.731861

**Authors:** Marie Horemans, Joren Stroobants, Joost Schepers, Marius Brusselmans, Bram Van Holm, Anne-Sophie Logist, Jelle Matthijnssens, Lieve Naesens, Kurt Vermeire, Guy Baele, Bert Vanmechelen

## Abstract

Nipah virus is a highly lethal, zoonotic paramyxovirus that has caused recurring outbreaks in several South and Southeast Asian countries since its discovery in Malaysia in 1998. Symptoms of infection include severe respiratory and neurological disease, often resulting in death. As no approved vaccines or antivirals are currently available to reduce the burden of this virus, it is classified as a biosafety level 4 pathogen. There is an urgent need for systems that enable research in a lower biocontainment setting, especially since the World Health Organization declared Nipah virus a priority pathogen for pandemic concern. In the past, several minigenome systems have already been developed as safe alternatives to working with infectious virus; however, these systems remain relatively inefficient and lack robustness and reliability for further applications. Therefore, we developed novel optimized RNA polymerase II-driven minigenomes with nanoluciferase or enhanced green fluorescent protein reporter genes. Both systems outperform previously designed Nipah virus minigenomes, are easily operable, and can be implemented for antiviral compound screenings.

## Introduction

Nipah virus (NiV) was first identified in Malaysia, during an outbreak among pigs and pig farmers from 1998 to 1999. A total of 265 human cases, including 105 deaths, were reported and over one million pigs were culled to prevent the virus from spreading (Chua et al., 2000). Since then, isolated cases and small outbreaks have occurred almost annually in South and Southeast Asian countries, primarily in India and Bangladesh (Li et al., 2023). Humans infected with NiV show a range of serious clinical manifestations, going from respiratory symptoms to severe neurological disease, often resulting in death: the case fatality rate (CFR) is estimated to range from 40% to 70%, but can be even higher depending on the outbreak (Devnath et al., 2022; Aditi and Shariff, 2019). Fruit bats of the genus *Pteropus* have been determined to be the natural reservoir of NiV (Li et al., 2023; Aditi and Shariff, 2019).

Two separate but closely related genetic lineages of NiV, with unique epidemiological and clinical features, can be distinguished: NiV-Malaysia (NiV-M) and NiV-Bangladesh (NiV-B). They share an overall nucleotide similarity of 92% and only differ six nucleotides in length: NiV-M is 18 246 nucleotides long, while NiV-B has a genome length of 18 252 nucleotides (Harcourt et al., 2005). During the Malaysian outbreak, spillover of NiV-M to the human population occurred through contact with infected pigs, with no observed human-to-human transmission (Aditi and Shariff, 2019). For NiV-B on the other hand, spillover to humans often occurs directly through bats without the involvement of an intermediary host. Additionally, NiV-B is associated with human-to-human transmission, more severe respiratory symptoms, and a higher CFR (Aditi and Shariff, 2019; Hauser et al., 2021).

Together with the zoonotic Hendra virus (HeV), NiV belongs to the *Henipavirus* genus within the *Paramyxoviridae* family. Paramyxoviruses are negative-sense, single-stranded, non-segmented RNA viruses that share a conserved genome organization of six genes (3’-N-P-M-F-G-L-5’), but this can be more depending on the genus (Rima et al., 2019). The N, P and L genes encode nucleocapsid-associated proteins: the nucleocapsid protein, the polymerase-associated phosphoprotein, and the large protein, including an RNA-dependent RNA polymerase, respectively. The remaining M, F, and G genes encode membrane-associated proteins: the matrix protein, fusion protein, and glycoprotein (Rima et al., 2019; Duprex and Dutch, 2023). A distinctive feature of paramyxoviruses is the “rule-of-six”, stating that genome lengths must be divisible by six in order to ensure proper encapsidation by the nucleocapsid protein, which in turn is necessary to enable efficient replication (Calain and Roux, 1993).

Due to its broad host range, high CFR, and the absence of effective vaccines and antivirals, NiV is classified as a biosafety level 4 (BSL-4) pathogen (Yoneda et al., 2006; Halpin and Rota, 2023). Consequently, this virus must be handled in high-containment facilities, significantly hindering research progress. However, safe alternatives to working with infectious virus, such as minigenome systems, have been developed to overcome this limitation. Minigenomes are truncated versions of the actual viral genome, consisting of a reporter gene flanked by the viral leader and trailer sequences. When cells are co-transfected with the minigenome and plasmids expressing the proteins of the viral replicase, reporter gene expression occurs. Minigenome systems are therefore safe to use in a BSL-2 environment, and can be implemented for the study of viral replication and transcription, or initial antiviral screenings (Haas and Lee, 2025).

The first NiV minigenome system, using a T7 promoter and a chloramphenicol acetyltransferase (CAT) reporter gene, was designed in 2003 by Halpin et al. (2004). A few years later, Freiberg et al. (2008) developed RNA polymerase (pol) I- and T7 pol-driven minigenomes, both using a CAT reporter gene. In 2024, Ke et al. (2024) published a T7 pol-driven system with nanoluciferase (Nluc) and green fluorescent protein (GFP) reporter genes, along with a cell line stably expressing N, P and L proteins instead of relying on plasmid-based expression. Finally, Wang et al. (2025) designed a T7 pol-driven system with firefly luciferase (Fluc) or enhanced green fluorescent protein (eGFP) reporter genes. Additional to minigenomes containing only a reporter gene, more complex systems have also been developed: Haas et al. (2024) designed a T7 pol-driven tetracistronic NiV minigenome, containing an mCherry reporter gene and NiV M, F and G genes. Besides transcription and replication, this system also enables budding, fusion and receptor binding.

These minigenomes have been proven valuable as tools for studying NiV, and certain improvements to their composition have been made over time. As CAT, Fluc or Nluc reporter genes have complex readout processes, relying on the addition of a substrate to produce measurable output, more straightforward reporter genes such as GFP or mCherry were applied. Contrary to the enzymatic reporters, GFP or mCherry directly exhibit fluorescence when expressed (Li et al., 2018). Using fluorescent reporter genes saves time as well as cost, two important factors when considering upscaling assays. However, the overall efficiency of minigenome assays remains limited, mostly due to T7 pol still being considered the gold standard as DNA-dependent RNA polymerase in minigenome design. T7 pol originates from the T7 bacteriophage, and is therefore not naturally present in mammalian cells (Borkotoky and Murali, 2018). As a result, it needs to be added exogenously through T7 expressing cell lines, or an additional plasmid. This complicates the transfection process and hereby limits the robustness and reliability for further usage of these systems.

Recently, the use of RNA pol II as an alternative to T7 pol as DNA-dependent RNA polymerase for reverse genetics has been proposed and successfully applied for other viruses (Nelson et al., 2017; Wang et al., 2015; Takahashi et al., 2023). Unlike T7 pol, RNA pol II is constitutively expressed in mammalian cells, hereby enhancing expression levels and removing the need for using exogenous T7 pol (Haas and Lee, 2024). We here present the first RNA pol II-driven minigenome system for NiV based on the original model of Halpin et al. (2004), using Nluc or eGFP reporter genes. These minigenomes demonstrate improved and robust reporter gene activity compared to previous models, are user-friendly and suitable for antiviral compound screening.

## Materials and methods

### Cell lines

Human lung adenocarcinoma epithelial cells (A549, CCL-185; ATCC, Manassas, VA, USA), baby hamster kidney cells stably expressing T7 polymerase (BSR-T7/5; gifted by Ian Goodfellow, University of Cambridge, UK), human embryonic kidney cells (HEK293T, CRL-3216; ATCC), and African green monkey kidney cells (Vero E6, CRL-1586; ATCC) were maintained in Dulbecco’s Modified Eagle Medium (DMEM; Thermo Fisher Scientific, Waltham, MA, USA) with 4.5 g/L D-Glucose and L-Glutamine, supplemented with 10% fetal bovine serum (FBS; Biowest, Nuaillé, France). 1% Penicillin-Streptomycin (Thermo Fisher Scientific) and 0.02% Gentamicin (50 mg/mL; Thermo Fisher Scientific) were added as antibiotics for each cell line. 0.2% Amphotericin B (250 µg/mL; Thermo Fisher Scientific) was added as antifungal for each cell line. 1% sodium bicarbonate (Thermo Fisher Scientific) was additionally added for A549, BSR-T7/5 and Vero E6 cell lines, and 0.5% Geneticin (100 mg/mL; InvivoGen, San Diego, CA, USA) was additionally added for BSR-T7/5 cells.

For use in antiviral compound screening, HEK293T cells were transduced with a lentiviral construct containing an mCherry reporter gene and blasticidin resistance gene using the ViraPower HiPerform T-Rex Gateway Expression System (Thermo Fisher Scientific). 7.5 µg/mL Polybrene Infection/Transfection Reagent (Merck kGaA, Darmstadt, Germany) was added to increase transduction efficiency. HEK293T medium supplemented with 10 µg/mL Blasticidin (Invivogen) was added for selection 48h after transduction, and was thereafter used for maintenance of HEK-mCherry cells.

### Plasmids

Synthetic DNA copies of NiV-B were ordered in five parts without any overlap through Twist Bioscience (South San Francisco, CA, USA). Support plasmids encoding NiV N, P and L (pCAGGS-N, pCAGGS-P and pCAGGS-L) were generated by PCR amplifying the respective genes from these synthetic sequences, using the Q5 Hot Start High-Fidelity 2X Master Mix (New England Biolabs, Ipswich, MA, USA) and by cloning these genes into a pCAGGS vector using the NEBuilder HiFi DNA Assembly Cloning Kit (New England Biolabs). As the C open reading frame (ORF) start codon falls within the P ORF, pCAGGS-P was afterwards mutated for the suppression of the expression of the C ORF through mutagenesis PCR (pCAGGS-Pmut). A plasmid encoding codon-optimized T7 polymerase (pCAGGS-T7opt) was a gift from Benhur Lee (Addgene plasmid # 65974 ; http://n2t.net/addgene:65974 ; RRID:Addgene 65974) (Addgene; Watertown, MA, USA) (Yun et al., 2015). Minigenome plasmids pT7-MG-Nluc, pT7-MG-eGFP, pCAGGS-MG-Nluc and pCAGGS-MG-eGFP were generated by cloning the eGFP gene or Nluc gene – flanked by the NiV leader and trailer regions – into a pT7 vector or pCAGGS vector, respectively. All minigenomes contain a hepatitis delta virus ribozyme (HDVRz) following the NiV leader sequence. pCAGGS minigenomes contain a hammerhead ribozyme (HHRz) between the CAG promoter and NiV trailer sequence. Additionally, minigenome plasmids pT7-HHRz-MG-Nluc and pCAGGS-noHHRz-MG-Nluc were generated by inserting an HHRz into pT7-MG-Nluc and by removing the HHRz from pCAGGS-MG-Nluc through PCR mutagenesis, respectively. All plasmids were transformed using NEB 5-alpha Competent *E. coli* (C2987H; New England Biolabs). Plasmid sequences were verified using nanopore sequencing (Rapid Barcoding Kit 96 V14 (SQK-RBK114.96) with Flongle flow cells (R10.4.1) (Oxford Nanopore Technologies, Oxford, UK)) and CLC Genomics Workbench (v22.0.3; QIAGEN, Hilden, Germany).

### Reverse transfection, Nano-Glo Luciferase readout and eGFP imaging

Reverse transfection was performed in HEK293T cells using Transporter 5 Transfection Reagent (Polysciences, Warrington, PA, USA) and in A549, BSR-T7/5 and Vero E6 cells using TransIT-LT1 Transfection Reagent (Mirus Bio, Madison, WI, USA) according to the manufacturer’s instructions. Reverse transfection was performed in a 96-well format. Per well, 8 ng pCAGGS-N, 2.5 ng pCAGGS-Pmut, 17.5 ng pCAGGS-L and 22 ng MG (for pCAGGS minigenomes) or 11 ng MG and 11 ng pCAGGS-T7opt (for pT7 minigenomes) was mixed with 5 µL 150 mM NaCl and 4:1 (DNA:reagent) Transporter 5 or 6.25 µL Opti-MEM (Thermo Fisher Scientific) and 3:1 TransIT-LT1, and incubated at room temperature for 20 minutes. To this mixture, 25 000 HEK293T cells or 10 000 A549, BSR-T7/5 or Vero E6 cells, resuspended in 50 µL medium, were added. Thereafter, the cell suspension with plasmid mixture was added to a 96-well plate prefilled with 100 µL medium. After 96h, Nano-Glo Luciferase readout or eGFP imaging was performed for Nluc or eGFP minigenomes, respectively. For Nano-Glo Luciferase readout, medium was removed and replaced with 50 µL phosphate-buffered saline (PBS). Per well, 1 µL Nano-Glo Luciferase Substrate was mixed with 50 µL Nano-Glo Luciferase Buffer (Promega, Madison, WI, USA) and added to the 96-well plate. Luminescence was measured within 3 to 120 minutes of adding the reagent using the GloMax Navigator Microplate Luminometer (Promega). For eGFP imaging, image acquisition was performed using the Operetta CLS High Content Imaging and Analysis system (Revvity, Inc., Waltham, MA, USA). High-resolution microscopic images were acquired using a 10X air objective (NA 0.3), capturing nine fields per well. For eGFP detection, excitation was achieved with a 460-490 nm LED, and fluorescence emission was collected using a 500-550 nm filter.

### Antiviral compound screening

Reverse transfection in HEK-mCherry cells using pCAGGS-MG-eGFP and required support plasmids was performed as described above. Cell suspension with plasmid mixture was added to a 96-well plate containing a 3-fold serial dilution of Remdesivir (GS-5734; MedChemExpress, Monmouth Junction, NJ, USA), starting at 30 µM. After 96h, image acquisition was performed using the Operetta High Content Imager (Revvity, Inc.). The image analysis algorithm was developed in-house using the Harmony^®^ software (Revvity, Inc., version 5.2). High-resolution microscopic images were acquired using a 10X air objective (NA 0.3), capturing nine fields per well. The mCherry fluorescent protein in the cell nucleus was excited using a 530–560 nm LED, and the corresponding emission was collected with a 570–650 nm filter. For eGFP detection, excitation was achieved with a 460–490 nm LED, and fluorescence emission was collected using a 500–550 nm filter. The image analysis algorithm was applied to quantify the total cell count and determine the number of infected cells. First, a basic flatfield correction was applied to the raw images, followed by background reduction in both the mCherry and eGFP channels. Afterwards, the intensity of the mCherry signal was used to identify and segment the cell nuclei. The eGFP signal was selected using an intensity threshold, and nonspecific signals were removed based on size. Finally, the percentage of infected cells was calculated as the ratio of infected cells to the total cell count. Infected cells were identified by the overlap of mCherry and eGFP signals, with every nucleus containing eGFP signal counted as infected. *Z*′ scores were calculated using the following formula: 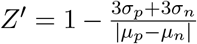 (*p* = positive control, percentage of infection of transfected cells; *n* = negative control, percentage of infection of untransfected cells) (Zhang et al., 1999).

### Statistical analysis

Both within-minigenome and between-minigenome differences were assessed using inverse-variance weighted one-way ANOVA. Within-minigenome differences compare absolute luciferase activity between support plasmid-transfected cells and negative control (NC) cells for each minigenome. Between-minigenome differences compare relative light units (RLUs) normalized to the average of each minigenome’s respective NC runs. Analysis of model residuals from a initial unweighted one-way ANOVA revealed substantial heteroscedasticity, supported by a positive Levene’s test. Average normalized RLU values were estimated using weighted least squares means. Significance tests for pairwise post-hoc comparisons of minigenomes were adjusted for multiple testing using Tukey’s procedure (Tukey, 1991). Significance level for all tests was set at *α* = 0.05. All statistical analyses were performed in CRAN R version 4.3.0 (R Core Team, 2021) using the `emmeans` (Lenth and Piaskowski, 2026) and `rstatix` (Kassambara, 2025) packages.

## Results

### RNA pol II-driven minigenome outperforms the original T7 pol-driven minigenome

Based on the original system of Halpin et al. (2004), we designed two minigenomes containing an Nluc reporter gene: a T7 pol-driven minigenome representing the original system as a control (pT7-MG-Nluc) and a novel RNA pol II-driven minigenome generated by replacing the T7 promoter with a CAG promoter and adding a self-cleaving HHRz downstream of the NiV trailer (pCAGGS-MG-Nluc) (Figure 1A). To compare the activities of both systems, HEK293T cells were transfected with either pT7-MG-Nluc or pCAGGS-MG-Nluc and required support plasmids; reporter gene expression was measured 96h post transfection. The L plasmid was omitted from the transfection as an NC. For pT7-MG-Nluc, a mean luciferase activity of 7.50 × 10^6^ RLU was measured, corresponding to a 29-fold increase in signal normalized to the NC (95% confidence interval (CI): [20 - 37]). Mean luciferase activity for pCAGGS-MG-Nluc was measured at 2.76 × 10^9^ RLU, corresponding to a 5 700-fold increase in signal normalized to the NC (95% CI: [5 350 - 6 050]) (Figure 1B & 1C). Therefore, our novel RNA pol II-driven system significantly outperforms the original system (*p ≤* 0.0001), demonstrating luciferase activity that is almost 200 times stronger.

**Figure 1:**
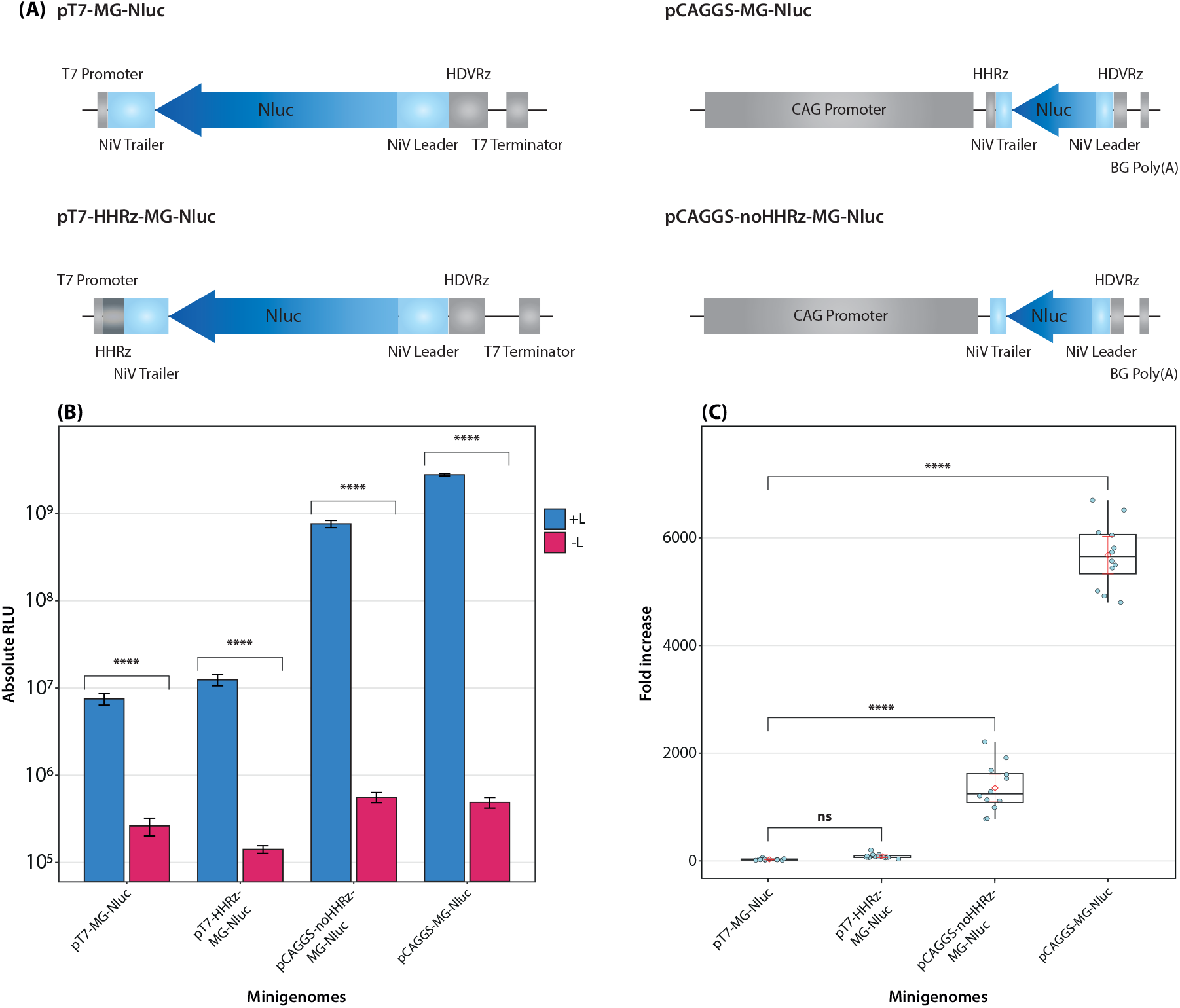
Comparison of composition and luciferase activity of all designed Nluc minigenomes. (A) Schematic representation of the designed minigenomes containing an Nluc reporter gene. HDVRz: Hepatitis delta virus ribozyme; HHRz: Hammerhead ribozyme; BG Poly(A): Betaglobin poly(A) signal. (B) HEK293T cells were transfected with minigenome plasmids and required support plasmids (+L); luciferase activity was measured 96h post transfection. The L plasmid was omitted from the transfection as an NC (-L). Data shown from two independent experiments with six replicates for each condition. Error bars indicate standard errors of the mean. A logarithmic scale is used for the y-axis. Statistical significance of differences between +L and -L values was assessed using an inverse-variance weighted ANOVA. Significance levels: **** = *p ≤* 0.0001. (C) Data from (B) shown as luciferase activity (+L) normalized to the NC (-L). Dots represent each luciferase measurement divided by the mean NC. Middle horizontal boxplot line represents the median, lower and upper horizontal lines represent the first and third quartile, respectively. Error bars in red show the estimated mean normalized RLU with a 95% CI. Statistical significance of differences in normalized luciferase activity between the original T7 pol-driven minigenome and other minigenomes was assessed using an inverse-variance weighted ANOVA. Significance levels: not significant (ns) = *p >* 0.05; **** = *p ≤* 0.0001.

To confirm that this gain in efficiency is attributable to the combined use of the CAG promoter and an HHRz, we additionally designed a T7 pol-driven system containing an HHRz (pT7-HHRz-MG-Nluc) and an RNA pol II-driven system lacking an HHRz (pCAGGS-noHHRz-MG-Nluc) (Figure 1A). pT7-HHRz-MG-Nluc showed only a limited increase in luciferase activity with a mean of 1.29 × 10^7^ RLU measured, corresponding to an 88-fold increase in signal normalized to the NC (95% CI: [62 - 113]), which was not significantly more than the fold increase observed in the original T7 pol-driven system (*p >* 0.05). Luciferase activity of pCAGGS-noHHRz-MG-Nluc was higher, but still far from exceeding that of pCAGGS-MG-Nluc, with a mean of 7.60 × 10^8^ RLU measured, corresponding to a 1 360-fold signal increase (95% CI: [1 100 1 620]) (Figure 1B & 1C). Compared to the original T7 pol-driven system however, this was a significantly higher fold increase in normalized luciferase activity (*p ≤* 0.0001). An important remark here is that these results cannot be attributed to different amounts of minigenome plasmid used for transfection, as varying these amounts did not greatly affect luciferase activity (Supplementary data).

Furthermore, we assessed the compatibility of pCAGGS-MG-Nluc in cell lines commonly used for NiV research. To this end, A549, BSR-T7/5, HEK293T and Vero E6 cells were transfected with pCAGGS-MG-Nluc and required support plasmids; reporter gene expression was measured 96h post transfection. The L plasmid was omitted from the transfection as an NC. The highest activity was observed in HEK293T cells, reaching a mean of 2.62 × 10^9^ RLU, corresponding to a 7 687-fold increase in signal normalized to the NC. In BSR-T7 and Vero E6 cells, the minigenome exhibited less, but still significantly higher mean luciferase activity, with 1.12 × 10^8^ RLU (246-fold increase in signal) and 2.28 × 10^8^ RLU (332-fold increase in signal) measured, respectively. No luciferase activity was measured in A549 cells (Figure 2).

**Figure 2:**
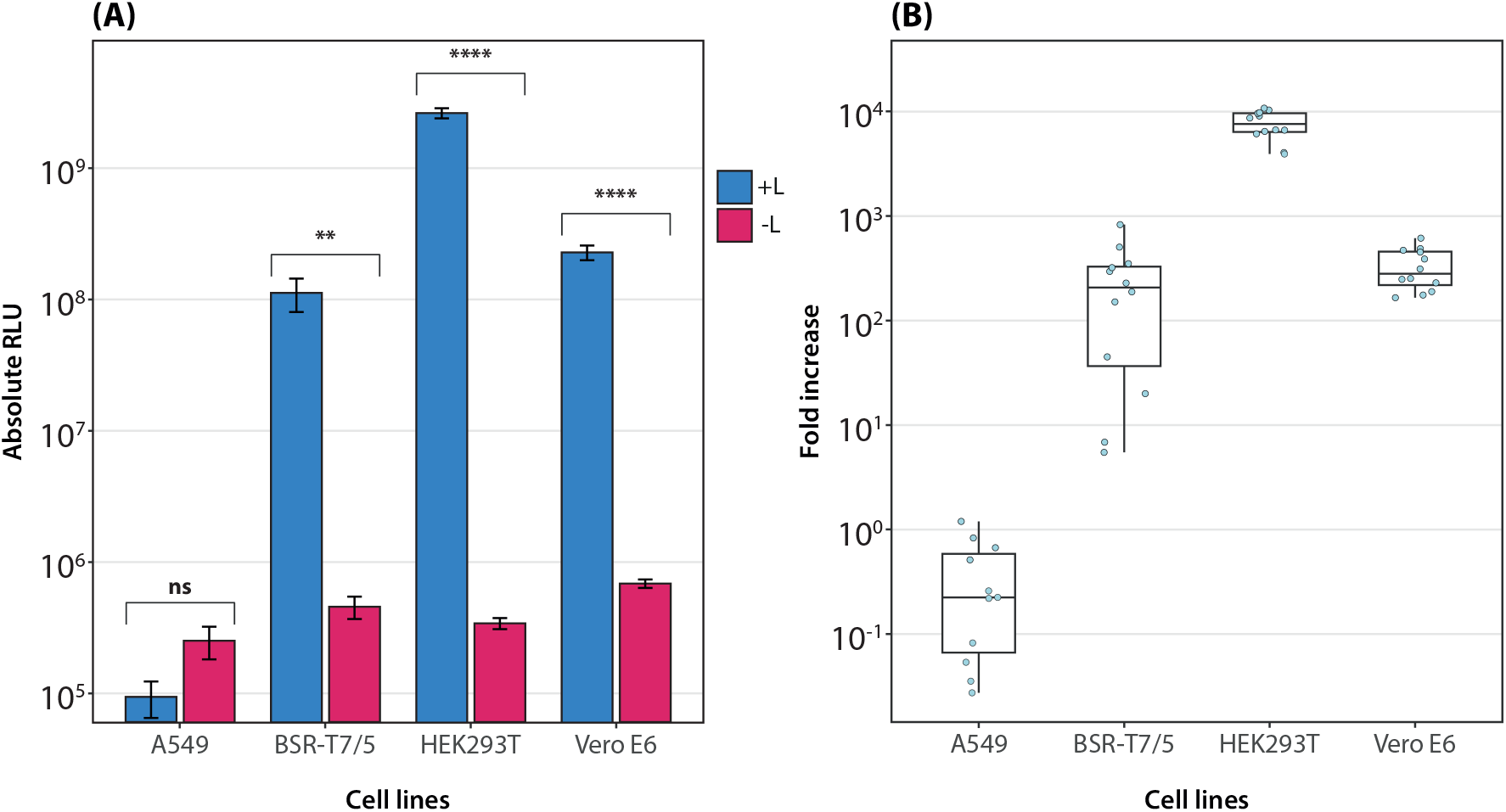
Compatibility of the RNA pol-II driven Nluc minigenome in cell lines commonly used for NiV research. (A) A549, BSR-T7/5, HEK293T and Vero E6 cells were transfected with pCAGGS-MG-Nluc and required support plasmids (+L); luciferase activity was measured 96h post transfection. The L plasmid was omitted from the transfection as an NC (-L). Data shown from two independent experiments with six replicates for each condition. One data point from the A549 +L group was omitted from the analysis due to a measurement error. Error bars indicate standard errors of the mean. A logarithmic scale is used for the y-axis. Statistical significance of differences between +L and -L values was assessed using an inverse-variance weighted ANOVA. Significance levels: ns = *p >* 0.05; ** = *p ≤* 0.01; **** = *p ≤* 0.0001. (B) Data from (A) shown as luciferase activity (+L) normalized to the NC (-L). Dots represent each luciferase measurement divided by the mean NC. Middle horizontal boxplot line represents the median, lower and upper horizontal lines represent the first and third quartile, respectively. A logarithmic scale is used for the y-axis.

### Development of an optimized eGFP minigenome system

To enable direct visualization of results without the need for adding reagents and to allow multi-day readouts, we designed an RNA pol II-driven minigenome containing an eGFP reporter gene (pCAGGS-MG-eGFP) based on pCAGGS-MG-Nluc (Figure 3A). HEK293T cells were transfected with pT7-MG-eGFP or pCAGGS-MG-eGFP and required support plasmids; eGFP readout was performed 96h post transfection. The L plasmid was omitted from the transfection as an NC. Activity of the RNA pol II-driven minigenome distinctly exceeded the activity of the T7 pol-driven system with mean eGFP SpotCounts of 1 358 and 11 per well, respectively (Figure 3B).

**Figure 3:**
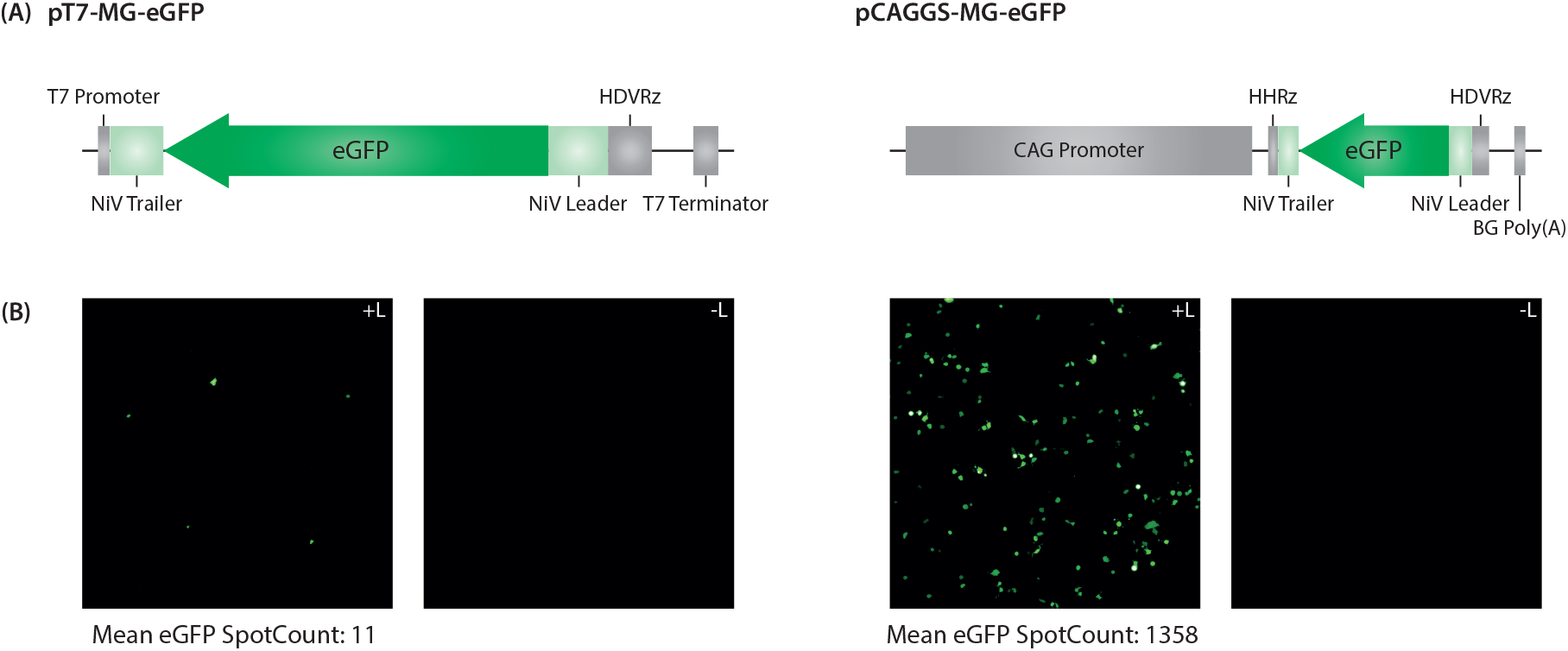
Comparison of eGFP minigenomes. (A) Schematic representation of the designed minigenomes containing an eGFP reporter gene. HDVRz: Hepatitis delta virus ribozyme; HHRz: Hammerhead ribozyme; BG Poly(A): Beta-globin poly(A) signal. (B) HEK293T cells were transfected with pT7-MG-eGFP or pCAGGS-MG-eGFP and required support plasmids (+L), images were taken 96h post transfection. The L plasmid was omitted from the transfection as an NC (-L). eGFP signal is shown in green. The number of eGFP spots per well was calculated over six replicates and is represented as the mean eGFP SpotCount.

### RNA pol II-driven eGFP minigenome as a tool for antiviral compound screening

Finally, we assessed the use of pCAGGS-MG-eGFP in an antiviral compound screening assay. For this, we developed HEK293T cells containing an mCherry reporter gene. This gene expresses a red fluorescent protein with a nuclear localization signal (NLS), enabling readout of the 50% cytotoxic concentration (CC50) of a compound. HEK-mCherry cells were transfected with pCAGGS-MG-eGFP and required support plasmids, and thereafter added to a 96-well plate containing a 3-fold serial dilution of Remdesivir. Readout was performed 96h post transfection. Remdesivir showed a dose-dependent inhibition of the minigenome system, with a 50% inhibitory concentration (IC50) of 0.304 µM and a CC50 of 1.713 µM (Figure 4A). We screened 19 additional compounds spread over a total of 10 experiments, but none of them showed minigenome inhibition. To assess the quality and robustness of the assay, *Z*′ scores of 10 independent experiments were calculated and returned values between 0.5 and 1.0 (Zhang et al., 1999), indicating a pronounced difference between the negative and positive controls, establishing that the assay is suited for high-throughput analysis (Figure 4B).

**Figure 4:**
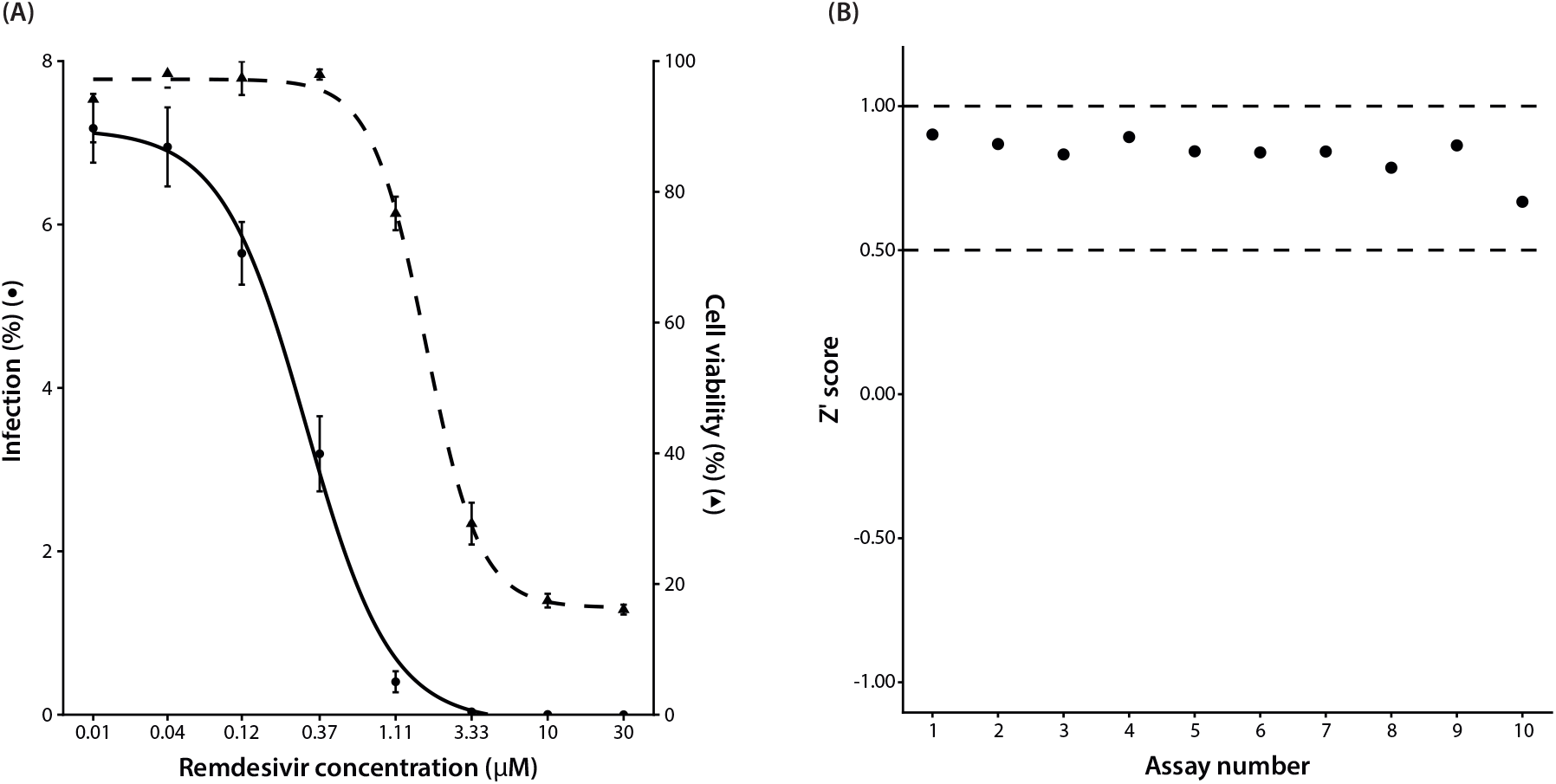
Use of the RNA pol II-driven eGFP minigenome for antiviral compound screening. (A) HEK-mCherry cells were transfected with pCAGGS-MG-eGFP and required support plasmids, and were thereafter added to a 96-well plate containing a 3-fold serial dilution of Remdesivir. eGFP readout was performed after 96h. Remdesivir shows a dose-dependent inhibition of the minigenome with an IC50 of 0.304 µM and a CC50 of 1.713 µM. Error bars indicate standard deviations. (B) *Z*′ scores of 10 assays were consistently situated between 0.5 and 1.0, indicating the robustness of the assay.

## Discussion

In the past, several NiV minigenome systems have been designed and successfully used to study different aspects of the virus, such as the functionality of C, V and W proteins or its compliance with the “rule-of-six” (Halpin et al., 2004; Sleeman et al., 2008). However, due to their reliance on exogenous T7 pol or complex readout processes, these systems remain relatively inefficient and lack robustness and reliability for further applications. Improved minigenome systems are urgently needed to accelerate the development of antivirals and to further investigate the NiV life cycle, as NiV is labeled a priority pathogen for pandemic concern by the World Health Organization (WHO) (Mallapaty, 2024). Based on the original minigenome by Halpin et al. (2004), we were able to design a novel, easily operable system that outperforms previously developed minigenomes.

The first described NiV minigenome system consists of a T7 promoter and a CAT reporter gene (Halpin et al., 2004). To enhance reporter gene expression, we replaced the T7 promoter with a CAG promoter, and the CAT reporter gene with an Nluc reporter gene. A CAG promoter is a composite promoter, consisting of the cytomegalovirus (CMV) enhancer, linked to the promoter and first intron of chicken beta-actin. This promoter is often used for its high-level expression, and additionally, it is recognized by the eukaryotic RNA pol II, hereby eliminating the need for adding exogenous T7 pol (Yew, 2005). However, a disadvantage of RNA pol II is the lack of a defined starting point for transcription. Since paramyxoviruses must comply to the “rule-of-six” for efficient replication, it is crucial that the minigenome has a length in nucleotides divisible by six. To solve this issue, we added a self-cleaving HHRz downstream of the NiV trailer. The first 10 nucleotides of this ribozyme are complementary to the trailer sequence, ensuring cleavage at the exact start of the trailer and a defined 5’ ending, thereby guaranteeing compliance to the “rule-of-six” (Figure 5). The importance of an HHRz for paramyxovirus reverse engineering systems has already been demonstrated by Beaty et al. (2017), who generated a T7 pol-based rescue system for NiV by incorporating an HHRz in their design, hereby enabling the use of the optimal T7 promoter sequence while still maintaining a genome length divisible by six. Additionally, Griffin et al. (2019) developed a NiV rescue system by combining the HHRz with a CMV immediate early promoter that is recognized by the eukaryotic RNA pol II.

**Figure 5:**
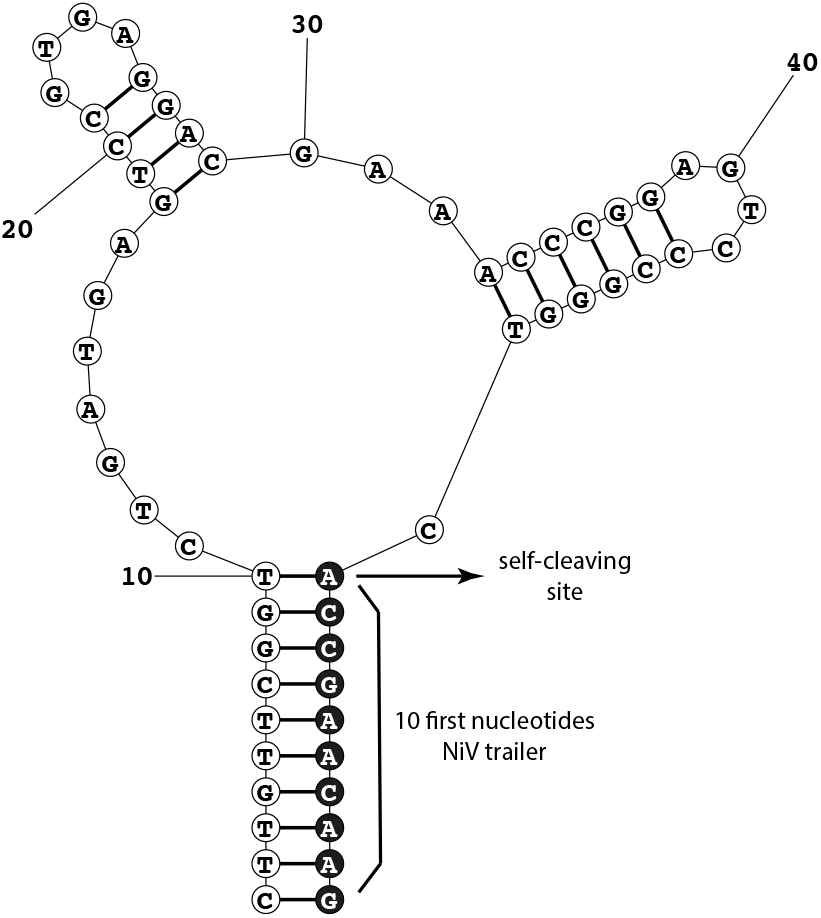
Self-cleaving hammerhead ribozyme (HHRz). Complementary nucleotides to the NiV trailer ensure cleavage at the exact start site of the minigenome, guaranteeing compliance to the “rule-of-six”. Figure generated using the MaxExpect Web Server from RNAstructure - Web servers for RNA Secondary Structure Prediction (https://rna.urmc.rochester.edu/RNAstructureWeb/Servers/MaxExpect/MaxExpect.html) and adapted using Adobe Illustrator.

We confirmed that the use of a CAG promoter together with an HHRz is optimal for minigenome design, yielding reporter gene activity nearly 200 times stronger compared to the original T7 pol-based system. Switching out the T7 promoter for a CAG promoter without incorporation of an HHRz produces a less pronounced, but still significant 45-fold increase in signal, highlighting the efficiency of the CAG promoter. In contrast, adding an HHRz to the T7 pol-driven system shows no significant fold increase in signal, which can be attributed to the fact that T7 pol has a defined transcription start site and does not need an HHRz for the minigenome to comply with the “rule-of-six”, even when using the optimal T7 promoter sequence.

Our optimal RNA pol II-driven minigenome is compatible with several cell lines commonly used in NiV research, and reaches its strongest activity in HEK293T cells. This provides a solid starting point for further research, e.g. antiviral compound screenings, as this is a human cell line. Less but still significant activity was observed for BSR-T7/5 and Vero E6 cells. Lastly, no minigenome activity was detected in A549 cells. It is however important to mention that we optimized the assay for HEK293T cells, and that adapted plasmid ratios or a different transfection reagent might result in a higher efficiency for other cell lines as well. Additionally, it should be taken into account that different cell lines demonstrate different transfection efficiencies.

To enable direct visualization of results without the need for adding substrate and to allow readouts over multiple days, we designed an RNA pol II-driven minigenome containing an eGFP reporter gene. The activity of this system in HEK293T cells is comparable to its Nluc counterpart with a mean eGFP signal more than 120-fold higher than that of the T7 pol-driven eGFP minigenome. Subsequently, we demonstrated that the RNA pol II-driven eGFP minigenome can be used for antiviral compound screening by assessing its susceptibility to Remdesivir. This is a nucleoside analog known to target the viral RNA polymerase of several RNA viruses, among which NiV (Bakheit et al., 2023). As expected, Remdesivir shows a dose-dependent inhibition of the minigenome in HEK293T cells with an IC50 of 0.304 µM. In comparison, the minigenome of Ke et al. (2024) shows susceptibility to Remdesivir in BHK21J cells expressing NiV N, P and L with an IC50 of 2.52 µM, while the minigenome of Wang et al. (2025) is inhibited with a half-maximal effective concentration (EC50) of 4.58 µM in A549 cells. Furthermore, the robustness and quality of this assay is indicated by Z’ values consistently situated between 0.5 and 1.

In conclusion, we designed novel RNA pol II-driven minigenomes containing Nluc or eGFP reporter genes. The combination of a CAG promoter with an HHRz provides an optimal composition for minigenome design, significantly outperforming the T7 pol-driven system. Furthermore, the eGFP minigenome enables direct visualization of results and multi-day readouts, and can be used for antiviral compound screening.

## Acknowledgements

The authors wish to thank dr. Piet Maes for his contributions during the initial stages of this study concerning funding and supervision. M.B. and G.B. acknowledge support from the DURABLE EU4Health project 02/2023-01/2027, which is co-funded by the European Union (call EU4H-2021-PJ4) under Grant Agreement No. 101102733. K.V. acknowledges the internal funding from the former Division of Virology and Chemotherapy, Rega Institute, KU Leuven. G.B. acknowledges support from the Research Foundation - Flanders (“Fonds voor Wetenschappelijk Onderzoek - Vlaanderen,” G098321N) and from the European Union Horizon 2023 RIA project LEAPS (grant agreement no. 101094685). B.V. acknowledges funding from the Belgian Federal Public Service Health, Food Chain Safety and Environment. A plasmid encoding codon-optimized T7 polymerase (pCAGGS-T7opt) was a gift from Benhur Lee (Addgene plasmid # 65974 ; http://n2t.net/addgene:65974 ; RRID:Addgene 65974) (Yun et al., 2015).

## Author contributions

Conceptualization of the study: B.V. Data curation: M.H., J.Sc., A.S.L. Formal analysis: M.H., J.Sc., M.B., G.B. Funding acquisition: J.M., K.V., G.B., B.V. Investigation: M.H., J.St., B.V.H., A.S.L., B.V. Methodology: J.St., B.V. Project administration: M.H., B.V. Resources: L.N., K.V. Supervision: J.M., G.B., B.V. Visualization: M.H. Writing - original draft: M.H. Writing -review and editing: all authors

