## Supplementary figures and images for "An optimized RNA polymerase II minigenome system for Nipah virus"

### Supplementary Data

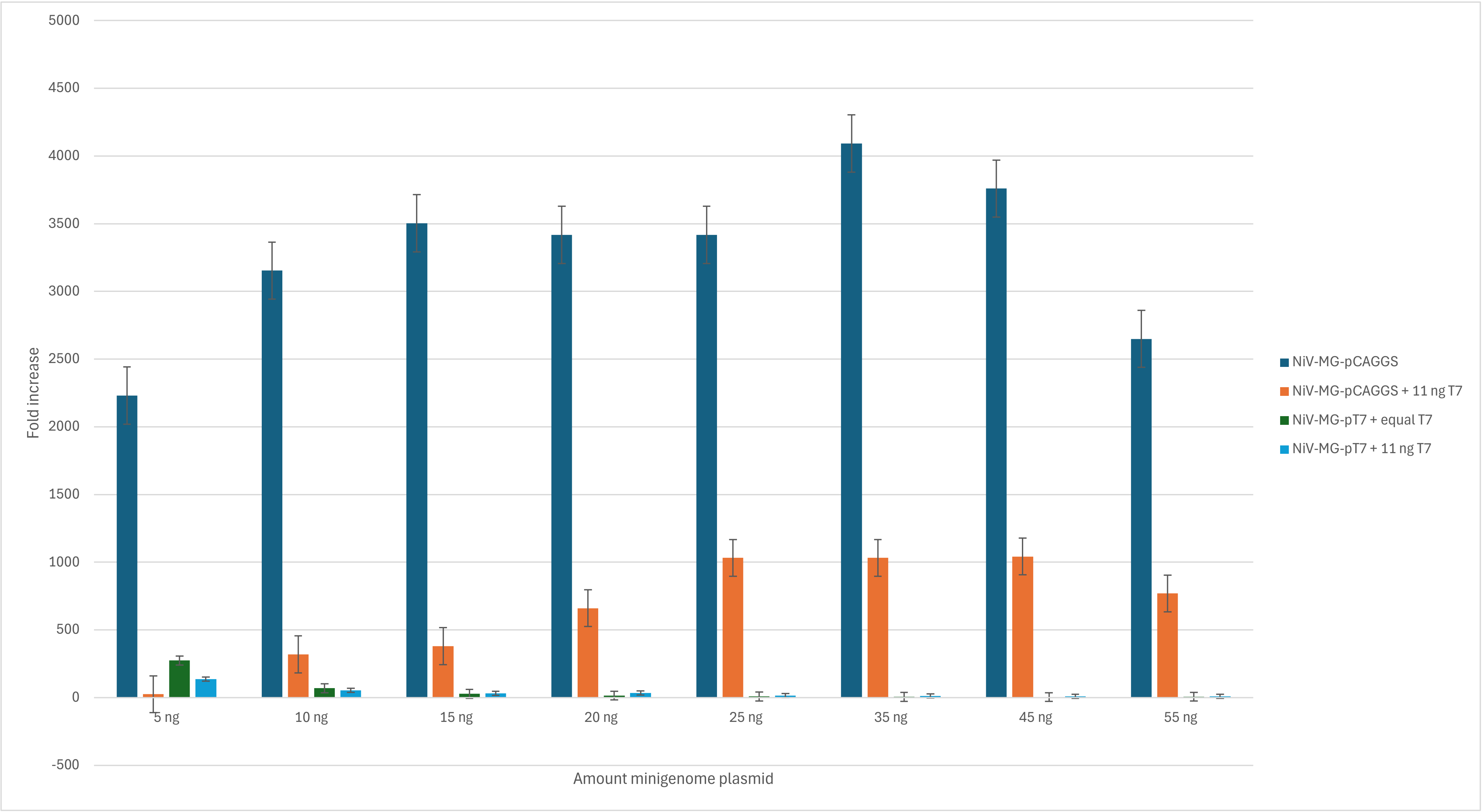
